# Should tissue structure suppress or amplify selection to minimize cancer risk?

**DOI:** 10.1101/062356

**Authors:** Laura Hindersin, Benjamin Werner, David Dingli, Arne Traulsen

## Abstract

**Background:** It has been frequently argued that tissues evolved to suppress the accumulation of growth enhancing cancer inducing mutations. A prominent example is the hierarchical structure of tissues with high cell turnover, where a small number of tissue specific stem cells produces a large number of specialised progeny during multiple differentiation steps. Another well known mechanism is the spatial organisation of stem cell populations and it is thought that this organisation suppresses fitness enhancing mutations. However, in small populations the suppression of advantageous mutations typically also implies an increased accumulation of deleterious mutations. Thus, it becomes an important question whether the suppression of potentially few advantageous mutations outweighs the combined effects of many deleterious mutations.

**Results:** We argue that the distribution of mutant fitness effects, e.g. the probability to hit a strong driver compared to many deleterious mutations, is crucial for the optimal organisation of a cancer suppressing tissue architecture and should be taken into account in arguments for the evolution of such tissues.

**Conclusion:** We show that for systems that are composed of few cells reflecting the typical organisation of a stem cell niche, amplification or suppression of selection can arise from subtle changes in the architecture. Moreover, we discuss special tissue structures that can suppress most types of non-neutral mutations simultaneously.

## I. BACKGROUND

It is a widely accepted view that tissues evolved to minimize the accumulation of somatic mutations during the live time of an individual [1–5]. Usually, this is achieved through the combined effect of multiple protective mechanisms. One important aspect is the hierarchical organization of most tissues, where few long lived stem cells give rise to a large and shorter lived population of progeny cells [6]. This allows for a high turnover of cells, while stem cells that are at risk to accumulate the potentially most harmful mutations, divide less frequently [1]. Thus most mutations occur at later stages of the hierarchy, where they are transient and are likely to be lost again due to the finite lifetime of most specialized cells [7]. The stem cell pool may exhibit additional layers of protection, including a slow rate of replication, membrane pumps to rapidly secrete genotoxic agents, elevated DNA repair mechanisms, and feedback loops to maintain certain spatial organizations [8, 9].

Another important contribution to the suppression of mutations might arise from particular ways of spatial stem cell organization [1, 2]. Here, we discuss properties of such spatially organized systems and how the actual realisation of the spatial organisation needs to take extrinsic risk into account, for example the actual distribution of mutant fitness effects.

Our theoretical results are based on the Moran process on graphs [10]. A population of cells is located on a graph, where the links of a focal cell indicate the neighboring cells that can be replaced by the offspring of the focal cell. New mutations have a relative fitness *r* > 0 compared to the wild-type with fitness 1 which influences their reproduction. One property of interest is the probability that a novel mutation takes over the whole population (reaches fixation on the graph). The graph represents the spatial structure of a population and is usually studied in comparison to a well-mixed population. Lieberman *et. al* defined a suppressor of selection as a graph that, compared to a well-mixed population, reduces the fixation probability of advantageous mutations (that have higher fitness than the wild-type) and increases the fixation probability of disadvantageous mutations (that have lower fitness than the wild-type) [10]. An amplifier is defined as the reverse: It increases the fixation probability of an advantageous mutation and decreases the fixation probability for a disadvantageous mutation, compared to the same mutation in a well-mixed population.

Often, it is implicitly assumed that suppressors of selection are desired, since they reduce the probability that a mutation that enhances the fitness of a cell reaches fixation within the stem cell population. This argument is at least partially the result of our limitation to reliably identify only strong drivers of selection in human malignancies. Although it has been known for a long time that some genomic alterations, such as mutations in the tumour suppressor genes TP53 and APC or the generation of fusion genes such as BCR-ABL are associated with specific tumors, often no known single driver mutation can be reliably identified. In these cases, either the driver oncogene is unknown, or the cancer phenotype is due to the combined effect of (many) mutations each with a small fitness effect. The current models of colorectal cancer development (adenoma to carcinoma sequence) or the progression of myelodysplastic syndromes to acute myeloid leukemia would be compatible with the latter model [11].

The properties of suppressors of selection have been the focus of research in several theoretical studies [1, 2, 12, 13]. A suppressor of selection can reduce the probability that a strong driver mutation reaches fixation from values close to 1 to 1 /*N*, where *N* is the size of the population at risk, e.g. the number of stem cells in a colonic crypt. However, classical examples of suppressors of selection come with a trade off, as they increase the probability of fixation of disadvantageous mutations from almost 0 to 1/*N*. As many mutations in evolutionary biology lead to a reduced fitness, this poses the question whether stem cell organization should ideally suppress or amplify selection. If most mutations are advantageous and thus lead to a growth advantage, a suppressor of selection would reduce the rate of evolution. However, if most mutations are disadvantageous, an amplifier of selection ensures that these mutant cells cannot take over the population. This would prevent the successive accumulation of many deleterious mutations within stem cell populations and minimize the risk of tissue failure such as aplastic anemia.

Taking these conflicting considerations into account leads to our main question of which tissue architecture and population dynamics are optimal for minimizing cancer risk. The answer to this question depends on the stem cell population size, the precise stem cell organization and the distribution of fitness effects of both single mutants and mutants further ahead in the path towards the full cancer phenotype. In the following, we address these points. In Section II we make an approximation for the rate of accumulation of mutations to compare the strongest suppressor of selection to the well-mixed population. Under the assumptions we make, the suppressor of selection reduces the accumulation of mutations if the total fraction of advantageous mutations is larger than 1/*N*. This first approximation leads to the question of how the distribution of fitness effects determines whether a suppressor or an amplifier of selection is useful to minimize cancer risk, which we study in section III. In Section IV, we study small graphs exemplifying the stem cell population at the base of the colonic crypt. Since cancer is usually caused by the accumulation of several mutations, in Section V we ask which kind of structure would be optimal to prevent the fixation of two consecutive mutations.

## II. FIXATION OF NOVEL MUTATIONS

We first consider a well-mixed population of size *N*, where a cell’s offspring can displace any other cell. In this case, the probability of fixation *ϕ*(*r*) of a single mutant cell that divides at rate *r* > 0 when non-mutated cells divide at rate 1 is [14, 15]

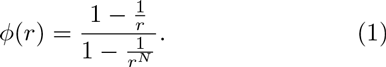

Throughout this work, we assume that the mutation rate is sufficiently small and the population size sufficiently large, such that we can consider one mutation at a time, i.e. we can neglect the effects of clonal interference [16–18]. Given a distribution of fitness effects *P*(*r*) and a mutation rate *μ*, the rate of accumulation of mutations in such a well-mixed population is

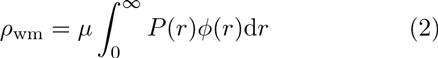

For simplicity, we first focus on large populations, *N* ≫ 1, where we have

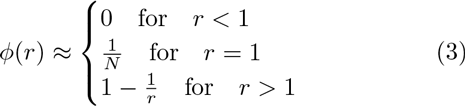

Dividing the distribution of fitness effects into *P*_<_(*r*) for disadvantageous mutations and *P*_>_(*r*) for advantageous mutations, we obtain for the rate of accumulation of mutations

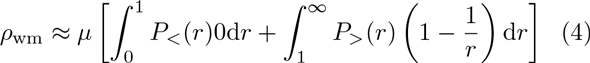

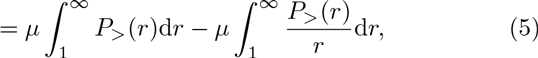

At most, the rate of accumulation of mutations is given by the mutation rate times the fraction of advantageous mutations, 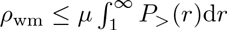.

On the other hand, consider a hypothetical population structure which could completely suppresses selection and leads to a neutral fixation probability which no longer depends on the selective advantage *r* [2, 10, 15]

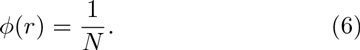

For this structure, the rate of accumulation of mutations is

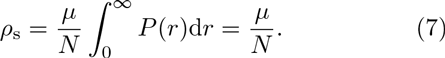

Note that here, all mutations reach fixation with the same probability. The extreme suppressor of selection leads to a reduced accumulation of mutations compared to the well-mixed case if

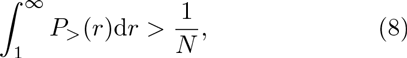

i.e. if the total fraction of advantageous mutations is larger than 1/*N*.

Alternatively, we can think of a population that amplifies selection, i.e. that leads to a higher probability of fixation for beneficial (advantageous) mutations and to a lower probability of fixation for deleterious (disadvantageous) mutations. In the approximation used above, we neglected the fixation of deleterious mutations, because their fixation probability is zero for large *N*. For an amplifier of selection, the second term in Eq. 5 becomes smaller and the rate of accumulation of mutations increases, as advantageous mutations will reach fixation with an even higher probability. Therefore the fraction of beneficial mutations must be smaller than in a well-mixed population to decrease the accumulation rate *ρ*.

On the other hand, taking into account deleterious mutations, an amplifier of selection can ensure that these reach fixation with an even lower probability compared to a well-mixed population. For large populations this probability is already small and the additional effect will not be of much further interest. But in small populations, it may be crucial to ensure that these mutations are weeded out. Such small populations can be of relevance in cancer initiation. For example, the colon is subdivided into many colonic crypts, each of them being maintained by a small independent subset of stem cells [19–21].

With these first arguments, it becomes clear that the fraction of beneficial and deleterious mutations is important for answering the question whether an amplifier or a suppressor of selection is the optimal structure to minimize cancer risk.

## III. THE DISTRIBUTION OF FITNESS EFFECTS OF CANCER MUTATIONS

In classical evolutionary biology, the distribution of fitness effects is of tremendous interest [22, 23]. The fitness effects of mutations depends on the evolutionary history of a population: If a population enters a novel environment, there can be mutations that lead to immediate fitness benefits. If a population has been evolving under constant conditions for a long time, it likely has already adapted to that specific environment and the chance that a novel mutation leads to beneficial effects decreases continuously. Usually, it is found that the vast majority of mutations are either deleterious or nearly neutral [24–26]. Only a small fraction of new mutations substantially increases evolutionary fitness.

Mutations that drive cancer initiation are usually thought to increase the fitness of a cell and only a few authors consider more general fitness landscapes [27]. Phenotypically these effects can be very diverse and include increased proliferation or decreased apoptosis rates, escape from an immune response or increased mutation rates [8, 21, 28, 29]. In most theoretical studies, advantageous mutations are considered to have small constant effects on fitness. In contrast, Durrett *et al.* have addressed the case of randomly distributed fitness of mutations in a branching process [30]. However, the authors have focused on growing populations, which seems more appropriate for the evolution of an already seeded tumor rather than the tumor initiation process within a healthy tissue.

If only advantageous mutations are dangerous in the sense that they can lead to cancer, an ideal tissue should suppress the accumulation of such mutations. A universal mechanism of protection is to reduce the effective mutation rate, e.g. by developing effective DNA repair mechanisms and proofreading by DNA polymerases. This is highlighted by the increased risk of cancer in patients with inherited defects in DNA repair mechanisms. However, even the best DNA polymerases cannot completely eliminate the risk of mistakes and a few mutations will always occur. Another mechanism is to kill (hyper) mutated cells. This is the task of TP53, highlighted by the increased risk of early malignancy in people with an inherited defect in TP53 (e.g. Li Fraumeni syndrome). Unfortunately, over-expressing TP53, or introducing multiple copies of TP53 in murine models leads to premature aging and actually, on average decreased life expectancy, although this may be a viable option in large, long-lived mammals such as elephants highlighting the enormous complexity and effects even of single genes [31]. There is the need for a balance between the mutation rate, DNA repair mechanisms and triggers of apoptosis within the cell to enable evolution of a species while reducing both the risk of cancer and early mortality. In addition, there might be alternative mechanisms that can suppress the spread of mutations within tissues. One such mechanism is the spatial organization of tissues.

Structures that suppress advantageous mutations usually also increase the fixation probability of mutations that cause reduced growth. In isolation, these mutations seem to be harmless because they are less fit than the wild-type cells. But they could interact with other subsequent mutations, leading to altered cell division properties via epistatic effects or environmental changes [27, 32, 33]. Such interactions between mutations have been investigated in experimental evolution in great detail [34–37], but they are usually neglected in the cancer community, partially because they are very difficult to measure. However, if initially disadvantageous mutations, which are arguably much more common, can turn to be dangerous for cancer initiation later, the organization of a tissue should adjust accordingly. An optimal tissue would in this scenario be an amplifier of selection, reducing the chances of fixation of the numerous disadvantageous mutations that can arise.

We study this by numerically calculating the fixation probability on an amplifier and suppressor graph of size 10. We use standard methods based on the transition matrix, which we generate from the adjacency matrix of the graph [13, 38, 39]. The transition matrix and the vector of fixation probabilities form a linear system of equations which can be solved for the fixation probabilities. To account for a broad range of fitness effects, we study mutations with fitness between 0 and 2, with 1 being the neutral reference fitness of the incumbent wild-type cells.

**FIG. 1.**
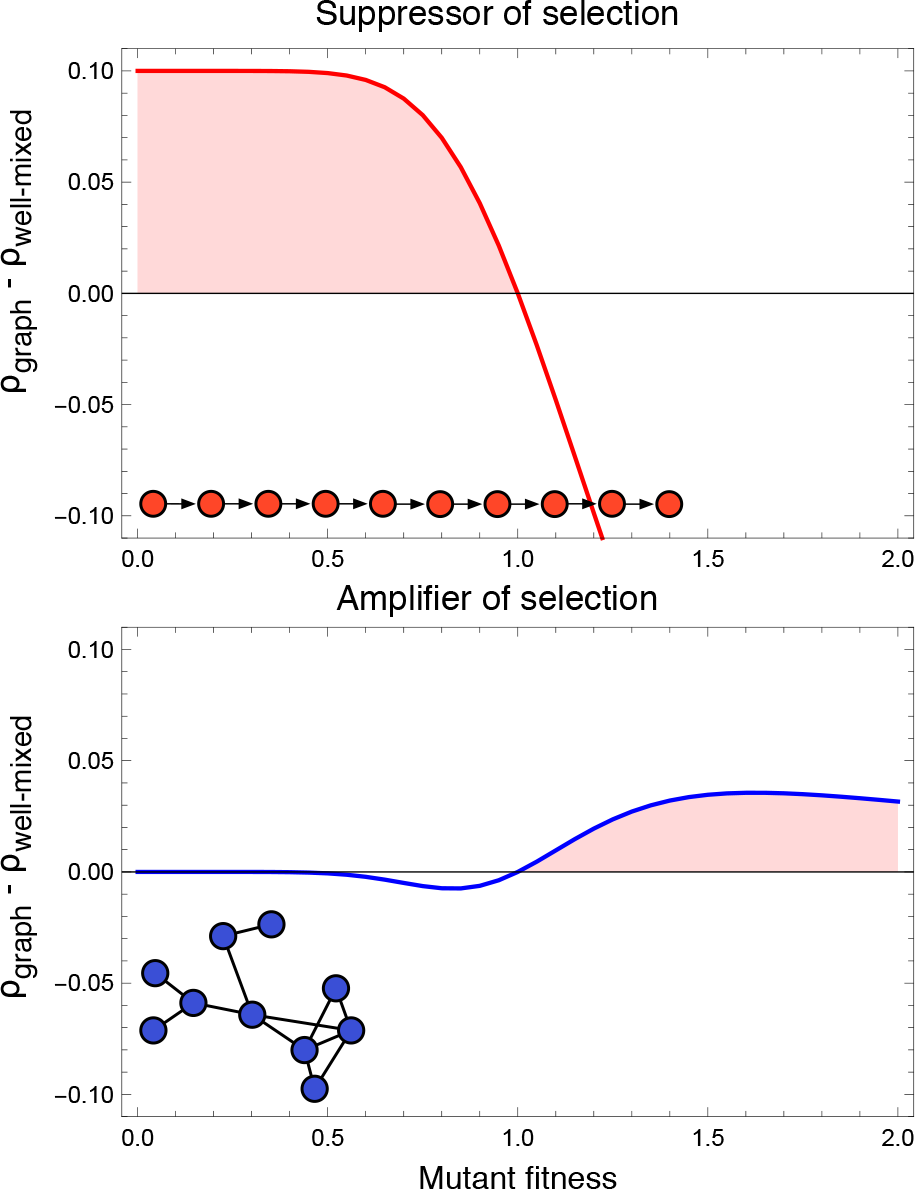
The difference between the fixation probability on a small graph and the fixation probability in an well-mixed population. **Top:** For a suppressor of selection, in this case the directed line, deleterious mutations have a higher fixation probability, whereas advantageous mutations have a lower fixation probability. The shaded region shows the fitness values for which mutations on this graph have a higher probability of taking over the population than in the well-mixed population. **Bottom:** For an amplifier of selection, here a random undirected graph, the fixation probability of advantageous mutations (shaded region) is increased, but the fixation probability of slightly deleterious mutations is decreased.

Fig. 1 illustrates these results for two small graphs, a suppressor and an amplifier of selection. We highlight the areas where mutations with this fitness effect on the corresponding graph have a higher fixation probability than in a well-mixed population of the same size. Depending on the distribution of fitness effects, the rate of accumulation of new mutations can be minimized by either an amplifier or a suppressor of selection. If most novel mutations are deleterious, a suppressor of selection could lead to faster accumulation of mutations. On the other hand, if most mutations lead to an immediate growth advantage, a tissue that suppresses selection could be better for minimizing the accumulation of cancer mutations. According to these arguments, the distribution of fitness effects becomes a crucial quantity in answering the question whether a suppressor or an amplifier of selection leads to a minimal cancer risk.

## IV. POPULATION STRUCTURES AND THEIR EFFECT ON FIXATION PROBABILITIES

Ideally, a tissue would decrease the fixation probability of both beneficial and deleterious mutations. Previously we have shown that graphs which suppress both beneficial and deleterious mutations can be constructed for some update mechanisms [13]. One example is the cycle which suppresses both beneficial and deleterious mutations. There is evidence suggesting that the stem cells of the intestinal crypts replace each other in a onedimensional way similar to neighbors on a cycle [4, 40]. However, the difference between the fixation probability on a cycle and a well-mixed population is relatively small and it remains unclear whether selection could lead to the evolution of such structures [41].

The simplest case of a structure that suppresses selection is a directed line [1]. The first cell is the root without incoming links and every cell can only replace its immediate successor. Mutations can only take over the whole graph if they occur in the root. Such a structure runs the risk of lineage extinction since there would be no redundancy in the system. One can envisage scenarios where this could lead to tissue failure and therefore place the organism at risk. It is perhaps for this reason that several stem cells occupy the base of each crypt in the colon.

To model the structure of colonic crypts, we consider small graphs that resemble this three-dimensional, bowl-like structure [20, 21]. The lowest layer of nodes corresponds to the stem cells in the bottom of the crypt. Links between nodes determine which cells can replace each other. With directed and weighted links, one can account for the outflowing cell dynamics by which the colonic epithelium is replenished. The properties of such a population structure depend strongly on the details of its implementation.

We consider two updating mechanisms:(i)Birth-death, where a cell is chosen for reproduction based on its fitness and replaces one of its neighbors with an identical copy of itself and (ii) death-Birth, where a random cell dies and its neighbors compete based on their fitness to replace the empty site with their offspring cell. Both update mechanisms can have different biological motivation [12, 42, 43]. It is still an ongoing discussion whether birth-death or death-birth updating is a more accurate description of cell dynamics in colonic crypts. In biology they are sometimes referred to as pushing or pulling, where the signal of proliferation comes either from the stem cells directly or is induced by feedback mechanisms from differentiated cells.

Graphs that suppress selection for both update mechanisms are rare, as most random undirected graphs are suppressors of selection for death-Birth updating, but amplifiers of selection for Birth-death updating [13]. Figure 2 shows four small graphs, which could represent the bottom part of a crypt, with different properties with respect to reducing or increasing fixation probabilities.

Figure 2a,b shows two examples of graphs which are suppressors of evolution for death-Birth updating [13]. Since all nodes have the same number of neighbors, mutants on these graphs have the same fixation probability as the well-mixed population for Birth-death updating (Isothermal Theorem, proven in [10]). However, these graphs reduce the fixation probability of both advantageous and deleterious mutations for death-Birth updating. Here, we neglect the walls of the crypt and only model the bottom by two rings. However, it seems reasonable to ignore the crypt walls, as there is an outwards replacement of cells [19] and the fixation of a mutation within the crypt bottom implies fixation within the whole crypt.

The bowl-like graph illustrated in Figure 2c suppresses selection for both Birth-death and death-Birth updating. Therefore, advantageous mutations have a lower fixation probability, but disadvantageous mutations have a higher fixation probability than the wild-type cells.

In Figure 2d, we implement directed and weighted links to model the outward replacement dynamics of the crypt bottom. The graph becomes an even stronger suppressor of selection compared to the same graph with undirected and equally weighted links.

All fixation probabilities have been calculated numerically based on the method described in [13, 39]. This approach is based on the numerical evaluation of the transition matrix of the Markov process associated with a graph, which naively scales with the graph size N as *2^N^ × 2^N^*. This allows us to obtain numerically exact results, but restricts the analysis to relatively small graphs (currently, for our implementation [39] up to 23 nodes).

These examples illustrate that it is far from obvious how tissues should be structured to prevent the accumulation of mutations.

## V. DOUBLE MUTATIONS

The initiation of cancer typically requires the accumulation of multiple mutations since a single mutated oncogene is rarely sufficient to cause cancer [8, 44, 45]. Thus, tissues architecture ultimately needs to prevent the accumulation of multiple mutations within single cells. Next, we study how a tissue should be structured to prevent the fixation of two consecutive mutations of different fitness effects. We assume that these mutations appear independently. Whether an amplifier or a suppressor of selection is more effective at preventing the accumulation of double mutations depends on the individual fitness effects of these mutations.

**FIG. 2.**
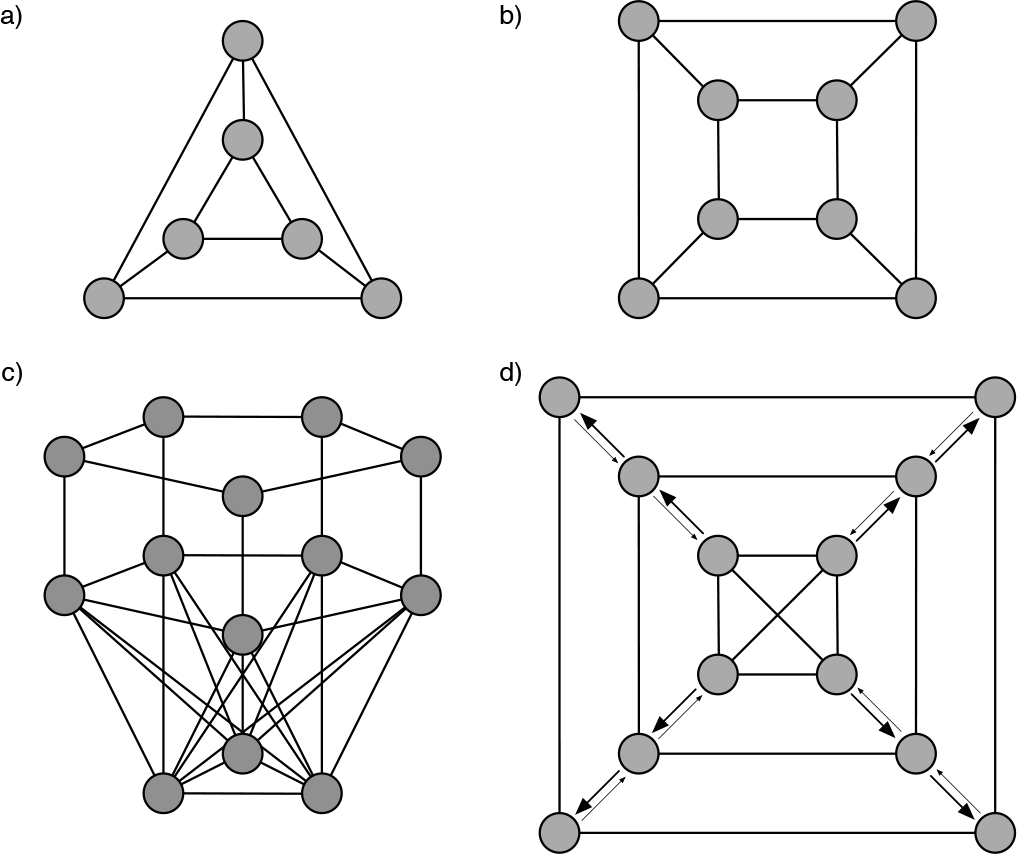
Different graph structures that can model colonic crypts. **a),b)**Two examples of suppressors of evolution for death-Birth updating. For Birth-death updating, these are equivalent to the well-mixed population in terms of the fixation probability. These two graphs consist of two layers of rings, their qualitative features do not change if we increase these structures to two rings of five, six, up to ten nodes. c) This bowl-like graph with 13 nodes comprised of a base of 3 interconnected nodes, which are all connected to all nodes of the middle layer of five nodes. From the middle layer, every node has a corresponding node in the upper ring, to which it is connected. The links are undirected and unweighted. This graph is a suppressor of selection for both Birth-death and death-Birth updating. Thus, it reduces the fixation probabilities for advantageous mutations. d)This graph has 12 nodes that are positioned in three layers. Here, the edges are directed and weighted. The outgoing edges between layers have a relative weight of 0.9, whereas the corresponding incoming weights are 0.1. This is to account for the outflowing cell-replacement of the colonic crypt. All other edge weights are 1. This graph is a suppressor of selection for Birth-death and death-Birth updating. The outflowing dynamics makes the suppression even stronger than in the same graph with unweighted edges

For example, consider the strongest possible suppressor of selection, e.g. the directed line. A mutated cell has a probability of 1/*N* to take over, independent of its fitness relative to the wild type cells. For simplicity, we focus on a single mutational path, i.e. we consider only a single order of mutations. The probability for two independent consecutive mutations is thus 1/*N*^2^. Let us compare this probability to that in a well-mixed population of the same population size. We study a system where the first mutation has relative fitness *r*_1_ and competes against the wild-type cells of fitness 1. The second mutation then has relative fitness *r*_2_ and competes in a population of cells with fitness *r*_1_. In this case, the combined fixation probability of the double mutant is

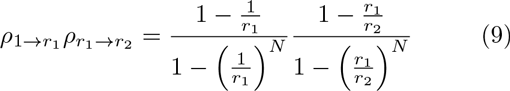

We now focus on mutations with small effects, *r*_1_ ≈ 1 and *r*_2_ ≈ *r*_1_, and we expand around *r*_1_ = *r*_2_ = 1,

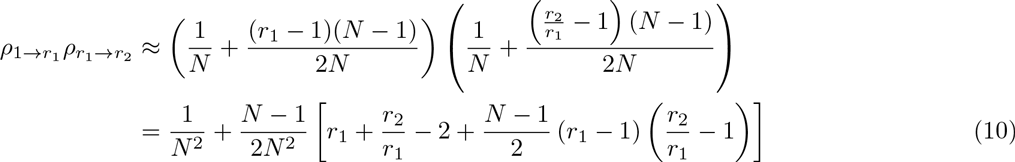

To see whether the directed line is more effective than the well-mixed population at preventing double mutations, we have to compare this to 1/*N*^2^. The directed line leads to a lower overall fixation probability if

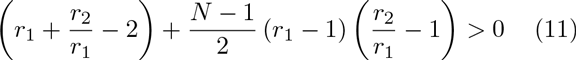

For two consecutive advantageous mutations, 1 < *r*_1_ < *r*_2_, this is clearly fulfilled and the directed line would reduce the fixation probability. In contrast, if the combined effect of two mutations is neutral (*r*_1_*r*_2_ = 1), but the first mutation confers either an advantage or a disadvantage, the fixation probability for the directed line is larger than the fixation probability of the well-mixed population, as the second, negative term always outweighs the first term for *N* > 3.

In general, the fixation probability of double mutants can either be larger or smaller on the directed line compared to the well-mixed populations. As illustrated in Figure 3, this directly depends on the choice of fitness values. In the shaded areas of Figure 3, the directed line is worse at preventing double mutations than the well-mixed population.

For many cases a strong suppressor of selection is worse at preventing the fixation of double mutations than the well-mixed population. If both mutations are advantageous, the directed line decreases the fixation probability. This effect prevails when one step is slightly disadvantageous, but as soon as the trajectory has to proceed through a sufficiently large fitness maximum or minimum, a well-mixed population performs better and suppresses such double mutants more efficiently. These results show that the term “suppressor of selection” can be misleading, because in some cases the “suppressor” actually accelerates the fixation of double mutations.

## VI. DISCUSSION

During homeostasis, stem cell replacement in the intestinal crypts is neutral [40]. Mutations that are commonly found in colorectal cancers likely give a competitive advantage to the cells [21]. This raises the question of how the tissue structure could act as a suppressor of novel mutations, decreasing the chance of advantageous mutations to take over the crypt.

Many graphs either act as a suppressor or amplifier of selection, preventing the fixation of either advantageous or disadvantageous mutations (compared to a well-mixed population), but not both [10, 12, 13, 46, 47].

However, these models have additional features that are not trivial: If mutations occur with a constant probability per division, then they do not arise with the same probability at all nodes. Instead, the arise with a probability proportional to the number of neighboring cells. In that case, a cycle-like structure with outflowing Birth-death cell replacement suppresses both beneficial and deleterious mutations [41].

The question of optimal tissue organization in order to minimize cancer risk is very complex. In general, the lifetime cancer risk is positively correlated with the number of stem cell divisions [48–50]. However, every tissue has unique needs and risks. As we have shown, subtle changes in tissue architecture can profoundly change its protective properties. This might explain why different tissues have evolved towards different organizations, but also why these organizations can be compromised by different types of mutations.

Furthermore, it is far from obvious that oncogenic mutations necessarily confer a direct fitness advantage to a cell. Deleterious mutations might play a role in cancer initiation via epistatic effects [27, 32]. Additionally, a mutated stem cell with lower replication rate could trigger its neighboring stem cells to compensate for the missing cell divisions by increased turnover and thus indirectly cause an increased cancer risk by effectively reducing the size of the active stem cell population.

**FIG. 3.**
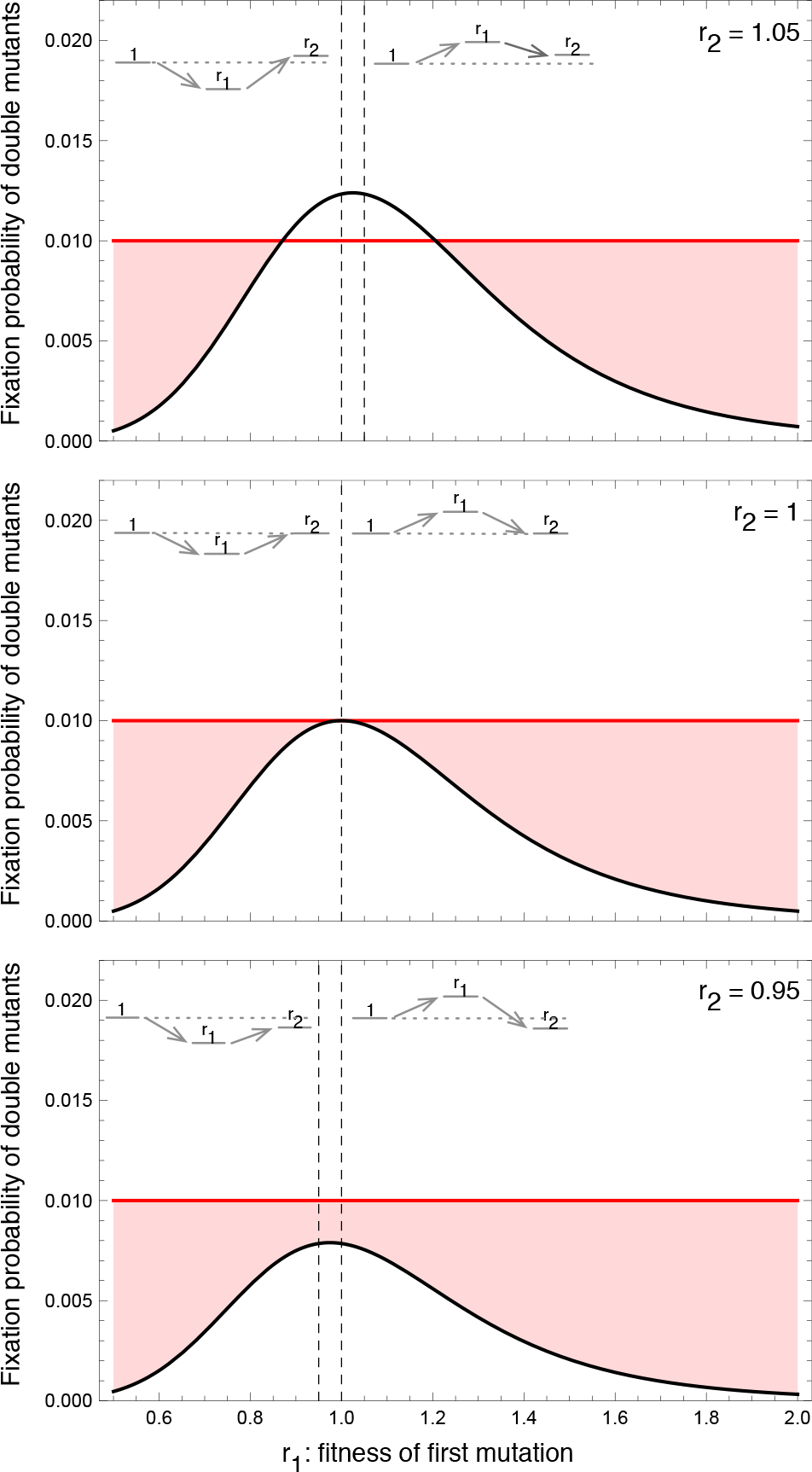
Fixation probability of two consecutive mutations on the directed line, a suppressor of selection (red), and the well-mixed population (black) of size *N* = 10. This fixation probability is plotted against the fitness of the first mutation. From top to bottom we show an advantageous second mutation (*r*_2_ = 1.05), a neutral (*r*_2_ = 1) and a disadvantageous (*r*_2_ = 0.95) second mutation. The shaded regions show the fitness values for which the fixation probability of the double mutations is higher on the directed line than in the well-mixed population.

Another risk is context dependent fitness, where the fitness of the mutant cell depends on the ecology (environment) of the cell population. For example, during development (growth) cell populations typically expand and favor mutant clones that grow faster, whereas in stationary conditions other phenotypes are selected for [33, 51, 52]. Thus, mutations that are neutral or disadvantageous at first, may become advantageous in later stages of development.

An additional effect of population structure is that the fixation of mutations becomes substantially slower in many graphs [18, 53], paving the way for increased clonal interference. It has been argued before that this will delay the accumulation of mutations and thus the onset of cancer [54].

We have shown that a “suppressor of selection” is not necessarily the optimal choice of a tissue structure in order to minimize cancer risk. It depends on the distribution of fitness effects. In order to prevent a sequence of two mutations, the directed line fares worse than the well-mixed population for most combinations of fitness values.

Overall, our approach is a step further to unravel the complexity of the spatial arrangement of a tissue to minimize the risk of cancer. A tissue architecture has to satisfy multiple requirements, some of which may appear to be conflicting. First and most importantly, it has to ensure the functionality of the organ. Secondly, it seems preferential to suppress harmful changes. Such changes can be manifold, and range from organ damage to cancer, normal or accelerated aging and possibly organ failure. We might have an intuitive understanding, why different organs evolved different architectures, and it seems natural that different architectures are prone to different errors. Nevertheless, our understanding of these problems is certainly incomplete and only recently has an evolutionary perspective become increasingly appreciated within the wider cancer research community. It is a hope that this might lead to a better understanding and ultimately a better approach towards cancer therapies.

## AUTHOR’S CONTRIBUTIONS

B.W., D.D., and A.T. have developed the concept. L.H. and A.T. have developed the analytical approach, L.H. has performed the numerical analysis. All authors have contributed to writing the manuscript. All authors have read and approved the manuscript.

## ACKNOWLEDGEMENTS

We thank Benjamin Allen, Andreas Deutsch and Ignacio Rodriguez-Brenes for constructive comments.

## References

[1] Nowak, M.A., Michor, F., Iwasa, Y.: The linear process of somatic evolution. Proceedings of the National Academy of Sciences USA 100, 14966–14969 (2003)

[2] Michor, F., Nowak, M.A., Frank, S.A., Iwasa, Y.: Stochastic elimination of cancer cells. Proceedings of the Royal Society B 270, 2017–2024 (2003)

[3] Komarova, N.L., Cheng, P.: Epithelial tissue architecture protects against cancer. Mathematical Biosciences 200, 90–117 (2006)

[4] Vermeulen, L., Morrissey, E., van der Heijden, M., Nicholson, A.M., Sottoriva, A., Buczacki, S., Kemp, R., Tavare, S., Winton, D.: Defining stem cell dynamics in models of intestinal tumor initiation. Science 342, 995–998 (2013)

[5] Bozic, I., Nowak, M.A.: Unwanted evolution. Science 342(6161), 938–939 (2013)

[6] Busch, K., Klapproth, K., Barile, M., Flossdorf, M., Holland-Letz, T., Schlenner, S.M., Reth, M., Hofer, T., Rodewald, H.-R.: Fundamental properties of unperturbed haematopoiesis from stem cells in vivo. Nature 518(7540), 542–546 (2015)

[7] Werner, B., Dingli, D., Lenaerts, T., Pacheco, J.M., Traulsen, A.: Dynamics of mutant cells in hierarchical organized tissues. PLoS Computational Biology 7, 1002290 (2011)

[8] Hanahan, D., Weinberg, R.A.: The hallmarks of cancer. Cell 100, 57–70 (2000)

[9] Hanahan, D., Weinberg, R.A.: Hallmarks of cancer: the next generation. Cell 144(5), 646–674 (2011)

[10] Lieberman, E., Hauert, C., Nowak, M.A.: Evolutionary dynamics on graphs. Nature 433, 312–316 (2005)

[11] Kandoth, C., McLellan, M.D., Vandin, F., Ye, K., Niu, B., Lu, C., Xie, M., Zhang, Q., McMichael, J.F., Wyczalkowski, M.A., Leiserson, M.D.M., Miller, C.A., Welch, J.S., Walter, M.J., Wendl, M.C., Ley, T.J., Wilson, R.K., Raphael, B.J., Ding, L.: Mutational landscape and significance across 12 major cancer types. Nature 502, 333–339 (2013). doi:10.1038/nature12634

[12] Kaveh, K., Komarova, N.L., Kohandel, M.: The duality of spatial death-birth and birth-death processes and limitations of the isothermal theorem. Journal of the Royal Society Open Science 2(140465) (2015)

[13] Hindersin, L., Traulsen, A.: Most undirected random graphs are amplifiers of selection for birth-death dynamics, but suppressors of selection for death-birth dynamics. PLoS Computational Biology 11, 1004437 (2015)

[14] Karlin, S., Taylor, H.M.A.: A First Course in Stochastic Processes, 2nd edition edn. Academic, London (1975)

[15] Nowak, M.A.: Evolutionary Dynamics. Harvard University Press, Cambridge MA (2006)

[16] Park, S.-C., Krug, J.: Clonal interference in large populations. Proceedings of the National Academy of Sciences USA 104(46), 18135–18140 (2007)

[17] Campos, P.R.A., Wahl., L.M.: The effects of population bottlenecks on clonal interference, and the adaptation effective population size. Evolution 63(4), 950–958 (2009)

[18] Frean, M., Rainey, P., Traulsen, A.: The effect of population structure on the rate of evolution. Proceedings of the Royal Society B 280, 20130211 (2013)

[19] Wright, N.A., Alison, M.: The Biology of Epithelial Cell Populations. Oxford University Press, USA, 198 Madison Avenue, New York, New York 10016 (1984)

[20] Baker, A.-M., Cereser, B., Melton, S., Fletcher, A.G., Rodriguez-Justo, M., Tadrous, P.J., Humphries, A., Elia, G., McDonald, S.A., Wright, N.A., Simons, B.D., Jansen, M., Graham, T.A.: Quantification of crypt and stem cell evolution in the normal and neoplastic human colon. Cell Reports 8(4), 940–947 (2014)

[21] Vermeulen, L., Snippert, H.J.: Stem cell dynamics in homeostasis and cancer of the intestine. Nature Reviews Cancer 14, 468–480 (2014)

[22] Eyre-Walker, A., Woolfit, A., Phelps, T.: The distribution of fitness effects of new deleterious amino acid mutations in humans. Genetics 173, 891–900 (2006)

[23] Eyre-Walker, A., Keightley, P.D.: The distribution of fitness effects of new mutations. Nature 8, 610–618 (2007)

[24] Zeyl, C., DeVisser, J.A.: Estimates of the rate and distribution of fitness effects of spontaneous mutation in Saccharomyces cerevisiae. Genetics 157, 53–61 (2001)

[25] Eyre-Walker, A., Keightley, P.D., Smith, N.G.C., Gaffney, D.: Quantifying the slightly deleterious mutation model of molecular evolution. Molecular Biology and Evolution 19, 2142–2149 (2002)

[26] Kassen, R., Bataillon, T.: Distribution of fitness effects among beneficial mutations before selection in experimental populations of bacteria. Nature Genetics 38, 484–488 (2006)

[27] Lipinski, K.A., Barber, L.J., Davies, M.N., Ashenden, M., Sottoriva, A., Gerlinger, M.: Cancer evolution and the limits of predictability in precision cancer medicine. Trends in Cancer 2(1), 49–63 (2016)

[28] Beerenwinkel, N., Antal, T., Dingli, D., Traulsen, A., Kinzler, K.W., Velculescu, V.E.E., Vogelstein, B., Nowak, M.A.: Genetic progression and the waiting time to cancer. PLoS Computational Biology 3, 225 (2007)

[29] Bozic, I., Antal, T., Ohtsuki, H., Carter, H., Kim, D., Chen, S., Karchin, R., Kinzler, K.W., Vogelstein, B., Nowak, M.A.: Accumulation of driver and passenger mutations during tumor progression. Proceedings of the National Academy of Sciences USA 107, 18545–18550 (2010)

[30] Durrett, R., Foo, J., Leder, K., Mayberry, J., Michor, F.: Evolutionary dynamics of tumor progression with random fitness values. Theoretical Population Biology 78(1), 54–66 (2010). doi:10.1016/j.tpb.2010.05.001

[31] Abegglen, L.M., Caulin, A.F., Chan, A., Lee, K., Robinson, R., Campbell, M.S., Kiso, W.K., Schmitt, D.L., Waddell, P.J., Bhaskara, S., Jensen, S.T., Maley, C.C., Schiffmann, J.D.: Potential mechanisms for cancer resistance in elephants and comparative cellular response to dna damage in humans. JAMA 314(17), 1850–1860 (2015)

[32] Bauer, B., Siebert, R., Traulsen, A.: Cancer initiation with epistatic interactions between driver and passenger mutations. Journal of Theoretical Biology 358, 52–60 (2014). doi:10.1016/j.jtbi.2014.05.018

[33] Werner, B., Traulsen, A., Dingli, D.: Ontogenic growth as the root of fundamental differences between childhood and adult cancer. Stem Cells 34, 543–550 (2016)

[34] Wolf, J.B., Brodie, E.D., Wade, M.J.: Epistasis and the Evolutionary Process. Oxford University Press, USA, 198 Madison Avenue, New York, New York 10016 (2000)

[35] Weinreich, D.M., Watson, R.A., Chao, L.: Perspective: sign epistasis and genetic constraint on evolutionary trajectories. Evolution 56(6), 1165–1174 (2005)

[36] Khan, A.I., Dinh, D.M., Schneider, D., Lenski, R.E., Cooper, T.F.:Negative epistasis between beneficial mutations in an evolving bacterial population. Science 332(6034), 1193–1196 (2011). doi:10.1126/science.1203801

[37] de Visser, J.A.G.M., Cooper, T.F., Elena, S.F.: The causes of epistasis. Proc Biol Sci 278(1725), 3617–24 (2011)

[38] Grinstead, C.M., Snell, J.L.: Introduction to Probability. American Mathematical Society, (1997)

[39] Hindersin, L., Möller, M., Traulsen, A., Bauer, B.: Exact numerical calculation of fixation probability and time on graphs. arXiv:1511.02696 0(0), 0 (2015)

[40] Lopez-Garcia, C., Klein, A.M., Simons, B.D., Winton, D.J.: Intestinal stem cell replacement follows a pattern of neutral drift. Science 330(6005), 822–825 (2010)

[41] Allen, B., Sample, C., Dementieva, Y., Medeiros, R.C., Paoletti, C., Nowak, M.A.:The molecular clock of neutral evolution can be accelerated or slowed by asymmetric spatial structure. PLoS Computational Biology 11(2), 1004108 (2015)

[42] Zukewich, J., Kurella, V., Doebeli, M., Hauert, C.: Consolidating birth-death and death-birth processes in structured populations. PLoS One 8(1), 54639 (2013)

[43] Débarre, F., Hauert, C., Doebeli, M.: Social evolution in structured populations. Nature Communications 5(3409) (2014)

[44] Fearon, E.R., Vogelstein, B.: A genetic model for colorectal tumorigenesis. Cell 61, 759–767 (1990)

[45] Vogelstein, B., Kinzler, K.W.: Cancer genes and the pathways they control. Nature Medicine 10, 789–799 (2004)

[46] Adlam, B., Nowak, M.A.: Universality of fixation probabilities in randomly structured populations. Scientific Reports 4(6692) (2014)

[47] Adlam, B., Chatterjee, K., Nowak, M.A.: Amplifiers of selection. Proceedings of the Royal Society A 471(2181), 20150114 (2015)

[48] Traulsen, A., Pacheco, J.M., Luzzatto, L., Dingli, D.: Somatic mutations and the hierarchy of hematopoiesis. BioEssays 32, 1003–1008 (2010)

[49] Tomasetti, C., Vogelstein, B.: Variation in cancer risk among tissues can be explained by the number of stem cell divisions. Science 347(6217), 78–81 (2015)

[50] Noble, R., Kaltz, O., Hochberg, M.E.: Peto’s paradox and human cancers. Philosophical Transactions of the Royal Society B 370(1673), 20150104 (2015)

[51] Rozhok, A.I., Salstrom, J.L., DeGregori, J.: Stochastic modeling reveals an evolutionary mechanism underlying elevated rates of childhood leukemia. Proceedings of the National Academy of Sciences 113, 1050–1055 (2016). doi:10.1073/pnas.1509333113

[52] Werner, B., Beier, F., Hummel, S., Balabanov, S., Las-say, L., Orlikowsky, T., Dingli, D., Bmmmendorf, T.H., Traulsen, A.: Reconstructing the in vivo dynamics of hematopoietic stem cells from telomere length distributions. eLife 4, 08687 (2016). doi:10.7554/eLife.08687

[53] Hindersin, L., Traulsen, A.: Counterintuitive properties of the fixation time in network-structured populations. Journal of The Royal Society Interface 11, 20140606 (2014)

[54] Martens, E.A., Kostadinov, R., Maley, C.C., Hallatschek, O.: Spatial structure increases the waiting time for cancer. New Journal of Physics 13 (2011). doi:10.1088/1367- 2630/13/11/115014

